# Trimeric Photosystem I facilitates energy transfer from phycobilisomes in *Synechocystis* sp. PCC 6803

**DOI:** 10.1101/2021.10.25.465751

**Authors:** Parveen Akhtar, Avratanu Biswas, Fanny Balog-Vig, Ildikó Domonkos, László Kovács, Petar H. Lambrev

**Author notes:** **Author for contact** Petar H. Lambrev. **Author Contributions** P.A. and P.H.L. conceived the project and designed the experiments. P.A., A.B., and F.B.-V. prepared the biological samples and conducted most of the spectroscopic measurements. L.K. and P.A. performed P_700_ oxidation measurements; I.D. and F.B.-V. did protein gel electrophoresis. P.A., A.B. and P.H.L. performed data analysis. P.A., A.B., and P.H.L. wrote the paper with contributions from all authors. P.H.L. is responsible for communication.

## Abstract

In cyanobacteria, phycobilisomes serve as peripheral light-harvesting complexes of the two photosystems, extending their antenna size and the wavelength range of photons available for photosynthesis. The abundance of phycobilisomes, the number of phycobiliproteins they contain, and their light-harvesting function are dynamically adjusted in response to the physiological conditions. Phycobilisomes are also thought to be involved in state transitions that maintain the excitation balance between the two photosystems. Unlike its eukaryotic counterpart, PSI is trimeric in many cyanobacterial species and the physiological significance of this is not well understood. Here we compared the composition and light-harvesting function of phycobilisomes in cells of *Synechocystis* sp. PCC 6803, which has primarily trimeric PSI, and the *ΔpsaL* mutant unable to form trimers. We also investigated a mutant additionally lacking the PsaJ and PsaF subunits of PSI. Both strains with monomeric PSI accumulated significantly more allophycocyanin per chlorophyll, indicating higher abundance of phycobilisomes. On the other hand, a higher phycocyanin:allophycocyanin ratio in WT suggests larger phycobilisomes or the presence of APC-less phycobilisomes (CpcL-type), that are not assembled in cells with monomeric PSI. Steady-state and time-resolved fluorescence spectroscopy at room temperature and 77 K revealed that PSII receives more energy from the phycobilisomes at the expense of PSI in cells with monomeric PSI, regardless of the presence of PsaF. Taken together, these results show that the oligomeric state of PSI has an impact on the excitation energy flow in *Synechocystis*.

**One-sentence summary:** Cyanobacterial mutants with monomeric PSI show changes in the composition and abundance of phycobilisomes and in the excitation energy transfer to PSII and PSI.

## INTRODUCTION

Although the photosynthetic apparatus of cyanobacteria is largely similar to that of eukaryotic algae and plants, cyanobacteria are distinct in several key aspects. Cyanobacteria lack the membrane-intrinsic antenna proteins of the LHC family but utilize membrane-peripheral phycobilisomes (PBSs) to increase the absorption cross-section of the two photosystems, PSI and PSII. The organization and function of the cyanobacterial thylakoid membranes is defined by the large PBSs attached to them and the coexistence of the photosynthetic and respiratory electron-transport chains sharing electron carriers. Apart from their peripheral antenna systems, cyanobacterial and plant PSI differ by their quaternary structure and subunit composition. Four small hydrophobic protein subunits - PsaF, PsaJ, PsaK and PsaX (only present in thermophilic species) cover the membrane-exposed surface of cyanobacterial PSI (Fromme et al., 2003). The functional role of PsaF in cyanobacterial PSI, apart from structural stabilization, is not established. It has been suggested that PsaF may be involved in the docking of the PBSs (Hippler et al., 1999) and, under iron deficiency, to mediate the interaction of the PSI core with the peripheral antenna IsiA (Fromme et al., 2003; Akita et al., 2020). Whereas eukaryotic PSI is monomeric, in cyanobacteria it is found in the form of trimers and in some species tetramers. The PsaL subunit is crucial for trimer formation and mutants lacking PsaL only accumulate PSI monomers (Chitnis and Chitnis, 1993). The physiological advantages of PSI oligomerization in cyanobacteria are not clear. PSI trimers have higher far-red absorption thanks to the presence of long-wavelength chlorophylls (Chls) and it may facilitate quenching of excess excitation energy by the oxidized reaction center (RC) and help protect against photoinhibition and ROS generation (Karapetyan et al., 1999; Kłodawska et al., 2020). It has also been proposed that the trimeric state could facilitate excitation energy transfer (EET) from the PBS to PSI (Şener et al., 2004).

The PBS is composed of phycobiliproteins (PBPs) and linker proteins organized as rods radiating from a membrane-attached core (MacColl, 1998; Arteni et al., 2009; Zheng et al., 2021). In the cyanobacterium *Synechocystis* sp. PCC 6803 (hereafter called *Synechocystis*), six PBS rods connect three hexameric phycocyanin (PC) discs each and the core consists of three cylinders, each with two stacked allophycocyanin (APC) hexamers (Arteni et al., 2009). The ApcD and ApcE (L_CM_) polypeptides of the core are crucial for the interaction with the photosystems and contain the longest-wavelength (680 nm) ‘terminal emitter’ pigments of the PBS that transfer energy to Chls in both photosystems (Ashby and Mullineaux, 1999; Rakhimberdieva et al., 2001; Liu and Blankenship, 2019). In situ cryoelectron tomography has revealed the ordered arrays of PBS-PSII supercomplexes, where energy can presumably migrate also laterally between PBS making for a very efficient light-harvesting system (Rast et al., 2019; Li et al., 2021). Plausible routes for energy migration from PBS to PSI can be directly via interaction between them (Mullineaux, 1994; Liu et al., 2013) or indirectly via “spillover” from PSII to PSI (McConnell et al., 2002; Ueno et al., 2017). ApcE is responsible for EET to PSII, whereas ApcD is proposed to serve as an energy donor primarily for PSI (Ashby and Mullineaux, 1999; Dong et al., 2009; Liu and Blankenship, 2019). In *Synechocystis* and other cyanobacterial species, an alternative PBS can be found, containing a single PC rod connected to the linker protein CpcL (CpcG2 in *Synechocystis*) but no APC core (Kondo et al., 2005; Mullineaux, 2008). The CpcL-type PBSs can interact with PSI transferring energy directly to it (Kondo et al., 2007; Watanabe et al., 2014).

The relative excitation of PSI and PSII can be rapidly regulated by the mechanism of state transitions, which is triggered by the redox state of the PQ pool (for reviews, see Mullineaux and Emlyn-Jones, 2005; Calzadilla and Kirilovsky, 2020). Several mutations in the PBS core are known to block or reduced the ability to perform state transitions, at least in some species (Ashby and Mullineaux, 1999; Dong et al., 2009; Calzadilla et al., 2019; Zlenko et al., 2019), highlighting the key role of the PBS in the process. However, the exact mechanism of cyanobacterial state transitions is under debate and alternative models are proposed, including a mobile PBS shuttling between PSI and PSII, regulated spillover (PSII–PSI energy transfer), and PSII quenching.

Cyanobacterial cells can modify the characteristics and abundance of PBSs in response to changes in the environmental conditions. Shortening of the PC rods under high growth light has been reported in several species, whereas low-intensity light elevates the PC content (Raps et al., 1985; Samson et al., 1994; Nomsawai et al., 1999). Macronutrient limitation results in extensive PBS breakdown (Salomon et al., 2013) that could provide the cell with amino acids – thereby, PBPs have obtained a secondary role as intracellular storage compounds.

Changes in the PBS can in turn affect the relative abundance of the membrane pigment-protein complexes. Genetically manipulated strains with shorter PBS rods or devoid of PBS accumulate more PSII relative to PSI (Nagarajan et al., 2014; Liberton et al., 2017). The PSI:PSII ratio in cyanobacteria is typically 2–4 and varies depending on the light intensity and quality that the organism is cultured in (Murakami and Fujita, 1991). Low growth irradiance increases the abundance of PSI and vice versa. The sensing mechanisms controlling the PBPs and photosystem abundance are not well elucidated (Stadnichuk et al., 2015). Different growth conditions, such as light, temperature and available nutrients may change the PSI oligomeric state and the ratio of monomers to trimers (Ivanov et al., 2006; Salomon and Keren, 2011; Kłodawska et al., 2015). Furthermore, mutants unable to form PSI trimers have shown changes in the relative fluorescence emission of PBS components suggesting that the oligomeric state influences the PBS composition (Kłodawska et al., 2020).

In this work, we take a look into the relationship between the oligomeric state of PSI, the PBS composition and abundance, and the EET from PBS to the photosystems, by comparing *Synechocystis* (which has predominantly trimeric PSI) and the Δ*psaL* mutant (unable to form trimers). In addition, we investigated a mutant lacking also the F and J subunits (ΔFIJL) to test if the PsaF and PsaJ subunits have specific role in EET from the PBS to PSI. We compare the PSI and PSII ratios as well as the PBP (PC and APC) content of the mutant cells with WT and will show that mutants with monomeric PSI have an altered composition and abundance of PBS. We also employed steady-state and time-resolved fluorescence spectroscopy at ambient and cryogenic temperature to evaluate the EET from PBS to the photosystems. By low-temperature time-resolved fluorescence spectroscopy we could separate PSII and PSI emission components and as well as populations of free, weakly, and strongly coupled PBPs. We show that the changes in the PBS composition affect the dynamics of EET in the cells and the excitation distribution between PSII and PSI, supporting the idea that PBS transfer energy more efficiently to trimeric than to monomeric PSI.

## MATERIALS AND METHODS

### Cyanobacterial cultures and preparations

A glucose-tolerant *Synechocystis* sp. PCC6803 strain, culturable under light-activated heterotrophic growth and maintained in our lab for decades, was used as WT. Cultures of WT, the Δ*psaL* mutant obtained on the same WT background (Kłodawska et al., 2015) and the subunit-depleted ΔFIJL mutant (Malavath et al., 2018) were grown photoautotrophically under continuous white fluorescent light (~35 μmol photons m^−2^ s^−1^) at 30°C. Thylakoid membranes and isolated PSI complexes were prepared as described in Akhtar et al. (2021).

PBSs were prepared according to the protocol described in Garnier et al. (1994) with some adjustments. Briefly, photoautotrophically grown cells were centrifuged to pellet at 6000 g at 25 °C. The pellet was washed twice with phosphate buffer (0.75 M phosphate buffer, 1 mM benzamidine hydrochloride hydrate, 1 mM EDTA, pH 7.0 and 1 mM of phenylmethylsulfonyl fluoride). The pellet was collected and treated with 0.2 % of lysozyme and incubated for 1 hour in dark at 37°C with continuous shaking at 200 rpm. After incubation cells were pelleted down by centrifugation at 6000 g for 7 minutes, 14 °C and washed twice in phosphate buffer to remove the remaining lysozyme. The cells were then broken with glass beads (≤ 106 μm diameter) using a homogenizer (Precellys Evolution) equipped with dry ice cooling compartment. The remaining glass beads were removed by centrifugation at 3000 g for 5 min at 14 °C. The supernatant was then treated with 3% Triton-X100 with continuous stirring for 30 min at room temperature in dark and centrifuged at 21,000 g for 30 min to remove the unsolubilized material. The appropriate sample fraction was collected and loaded onto a sucrose density step gradient (0.25, 0.5, 0.75, 1 M) and centrifuged for 16 h at 104,000 g 14 °C for further purification. The gradient fraction containing PBSs was collected.

### Pigment analysis

Chls were extracted from the cell suspensions in 90% methanol and the Chl contents were determined spectrophotometrically using molar absorption coefficients described in Lichtenthaler (1987).

The phycobiliprotein content was determined as described in Zavřel et al. (2018). For PBPs isolation, 100 ml of the cells from each type were pelleted by centrifugation at 6000 g for 5 min and resuspend in phosphate buffer (50 mM, pH 6.5) to total vol of 5 ml. The cells were broken using a Precellys Evolution homogenizer with dry-ice-cooled chamber (10 cycles of braking, 30 s vortex, 5500 rpm and two min cooling). The homogenate was then sonicated intermittently (five sec sonication with interval of 10 sec rest, four times) by ultrasonicator at ice water temperature. Unbroken cells and cell debris were removed by low-speed centrifugation. Cell homogenates were then ultracentrifuged at 104 000g for 60 min. Absorption spectra of the transparent supernatant in the range of 220–750 nm were recorded to determine the soluble PBP content of the cells.

### Quantification of PSI and PSII

The PSI and PSII concentrations were determined spectrophotometrically using the protocol described by Fujita and Murakami (1987). For P_700_ measurements the samples were suspended to 20 μg/ml (or optical density of 2 at 680 nm) and for cytochrome (Cyt) to 60 μg/ml (or optical density of 6 at 680 nm) in a buffer containing 20 mM MES/NaOH, pH 6.4, 10 mM MgCl_2_, 10 mM CaCl_2_. The control absorption spectra in the range of 350–750 nm were recorded from each sample. To estimate the concentration of P_700_ and Cyts, absorption spectra were recorded in the range 650–750 and 500–600 nm, respectively, with bandwidth of 1 nm. The P_700_ was first oxidized with 1 mM potassium ferricyanide and then reduced with sodium ascorbate. The difference spectra (690–720 nm) between oxidized and reduced P_700_ were identical to P_700_ and the difference at 700 nm was taken as the signal of P_700_. P_700_ abundance was estimated from the absorption difference with a molar extinct coefficient Δε_ox-red_ = 64 mM^−1^ cm^−1^ at 700 nm. For PSII determination, first all Cyts were oxidized with 1 mM of potassium ferricyanide. Then few grains of hydroquinone were added, followed by addition of sodium ascorbate and sodium dithionite. The difference spectra (520–580 nm) between hydroquinone-reduced and ascorbate-reduced had peak at 559 nm – Cyt *b*_559_.

### Redox kinetics of P700

The functional antenna size of PSI was estimated by the rate of light induced oxidation of P_700_ RC under light limiting conditions. The oxidation kinetics of P_700_ upon illumination was followed by the measurement of absorbance change at 830 nm using Dual-PAM 100 Chl a fluorometer (Walz, Germany). Prior to measurement samples were dark-adapted for 3 minutes in the presence of 100 μM methylviologen (MV) and 20 μM DCMU (3-(3,4-dichlorophenyl)-1,1-dimethylurea) then cell suspension equivalent to 20 μg Chl was filtered onto a 25 mm diameter glass fibre syringe filter disc (Whatman GF/C). The filter discs, placed between two microscopy slides with a spacer, were inserted between the fibre optics of the emitter-detector unit. Samples were illuminated with 5-s long 635-nm pulses at various intensities (6, 31, 140, 251, and 805 μmol photon m^−2^ s^−1^) consecutively and the oxidation kinetics were recorded at a millisecond sampling rate.

### Steady-state absorption and fluorescence spectroscopy

Absorption spectra in the range of 350–750 nm were recorded at room temperature with a Thermo Evolution 500 dual-beam spectrophotometer. The measurements were performed in a standard glass cell of 1-cm optical path length with 1 nm spectral bandwidth.

Fluorescence emission spectra in the visible range were measured from the same samples at room temperature and 77K on a FP-8500 (Jasco, Japan) spectrofluorometer. The sample were diluted to absorbance of 0.1 per cm at the red maximum. Emission spectra in the range of 620–780 nm were recorded with excitation wavelength of 440 nm and 580 nm and excitation/emission bandwidth of 3 nm. The measurements were performed with 1 nm increment and 1 s integration time. For measurements at 77 K, samples were cooled in an optical cryostat (Optistat DN, Oxford Instruments, UK). The spectra are corrected for the spectral response of the detector.

### Time-resolved fluorescence spectroscopy

Picosecond time-resolved fluorescence measurements were performed with a time-correlated single-photon counting instrument (FluoTime 200/PicoHarp 300 spectrometer, PicoQuant). Excitation was provided by Fianium WhiteLase Micro (NKT Photonics, UK) supercontinuum laser, generating white-light pulses with a repetition rate of 20 MHz. Excitation wavelengths of 440 and 580 nm were used to excite selectively Chls and PBSs. The fluorescence decays were recorded at wavelengths of 600–744 nm with 8 nm steps, at room temperature, and 605–760 nm with 5 nm steps at 77 K. All the samples were diluted to an absorbance of 0.03 at excitation wavelength. For the room temperature measurements, the suspension (whole cells or isolated complexes) was placed in 1 mm flow cell and circulated at a flow rate of 4 ml/min. For 77 K measurements, the suspension was placed in a 1 mm demountable cryogenic quartz cell and cooled in an optical cryostat (Optistat DN, Oxford Instruments, UK). The total instrument response (IRF) measured using 1% Ludox as scattering solution has width of 40 ps. The data are corrected for the spectral response of the detector. Global multiexponential lifetime analysis with IRF reconvolution was performed using MATLAB.

### Statistical analysis

Whenever appropriate, data are presented as mean ± standard error, obtained from independent measurements on different cell batches. The statistical significance, or lack thereof, of differences between the two mutant strains and the WT is reported based on Student’s *t*-test (*p* < 0.05).

## RESULTS

### Changes in the pigment stoichiometry

We cultured *Synechocystis* (WT), which contains PSI trimers and the two mutants with monomeric PSI, Δ*psaL* and ΔFIJL, under the exact same conditions, to examine the phenotype effects of the mutations. Both mutants appeared more greenish in colour suggesting a change in the pigment composition of the cells. Accordingly, absorption spectra of the supernatant obtained after sedimenting the broken cell debris (Supplementary Fig. S1) show a distinct shoulder around 650 nm in both mutants – Δ*psaL* and ΔFIJL – suggesting increased APC content. We estimated the PC and APC composition of the cell cultures from absorption spectra of the supernatant (Table 1). The Chl content was measured from methanol extracts of either the cell debris or whole cell sediment, yielding approximately equal results. We found that the ratio of PC to Chl was unchanged between the WT and mutant cultures. However, the amount of APC, on a PC or Chl basis, was significantly higher in the monomeric mutants – up to 75% more than in the WT, as shown by the PC:APC and Chl:APC ratios. If we assume that all, or almost all, PC in the WT is found in the PBS rods, the lower PC:APC ratio means that the PBS in the mutants has either fewer or shorter PC rods. On the other hand, the more abundant APC indicates a higher number of PBS cores in the cells. From these data it follows that both the composition and the number of the PBSs are altered in the mutants with monomeric PSI – they contain more PBSs as a whole (on Chl basis) compared to WT, but the PBSs in WT are larger, containing more PC, or WT cells contain additional PBSs without APC (see below).

**Table 1.**
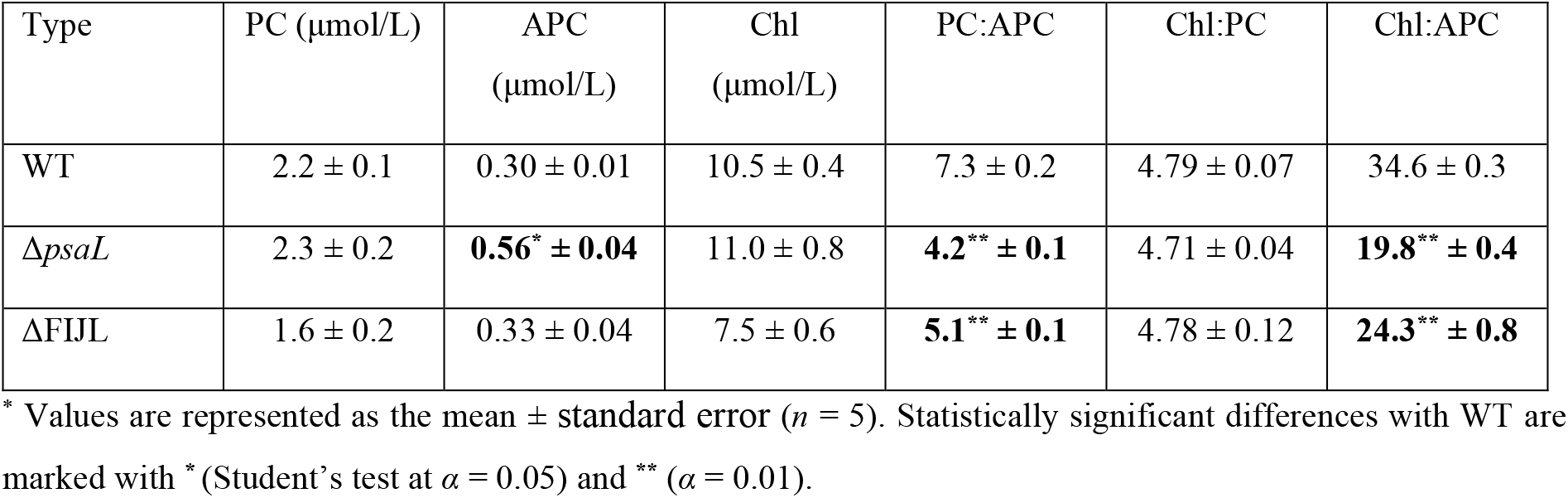
Pigment content of cell cultures

It must be noted that the PC:APC ratios in the WT are higher than expected. PBSs are typically found to contain six rods with three PC hexamers per tricylindrical core (Arteni et al., 2009; Rast et al., 2019), which amounts to a PC:APC ratio of 3:1. Although one could merely attribute the discrepancy to a systematic error in the PC:APC estimation, it should be pointed out that we obtained different results from isolated PBSs using the same measurement methodology (Supplementary Table S1). The PC:APC ratio of isolated PBSs was found to be around 4:1 – similar to previous reports. More importantly, the ratios were the same in PBSs of the WT and mutants, as evident from their nearly identical absorption spectra (Supplementary Fig. S1). It must be concluded that WT cells under our growth conditions contain extra PC, which is either not connected to the PBS core in vivo or is weakly connected such that it dissociates during isolation of PBSs.

The striking change in the PBS content and composition found in the monomeric PSI mutants compared to WT *Synechocystis* prompted to test whether there was a corresponding change in the photosystem stoichiometry. To this end, we estimated the number of PSI and PSII RCs in the cells by the absorption difference of the oxidized and reduced forms of P_700_ and Cyt *b*_559_, respectively (Table 2 and Supplementary Fig. S2). We found P_700_:Cyt *b*_559_ ratios of 1.5–1.7 that were not significantly different between mutants and WT. These values are similar to the ones reported by Murakami and Fujita (1991). However, in this paper it was assumed that there are two Cyt *b*_559_ per PSII RC, hence the reported PSI:PSII ratios were twice as high. A RC ratio of 1.5 means that there are equimolar ratios of PSI trimers (PSI_3_) and PSII dimers (PSII_2_) in WT or three monomeric PSI per PSII_2_ in the mutants. From these ratios and the number of Chls in PSI and PSII, we can estimate 3–4 PSI and the same number of PSII complexes per PBS in the WT. In the monomeric mutants, although the number of PBS is apparently increased, there are still more (monomeric) PSI complexes per PBS (6–7) but fewer PSII (2–3). Thus, we could potentially interpret the increased number of PBS in the mutants as an adaptive response that compensates for the number of PSI per PBS.

**Table 2.**
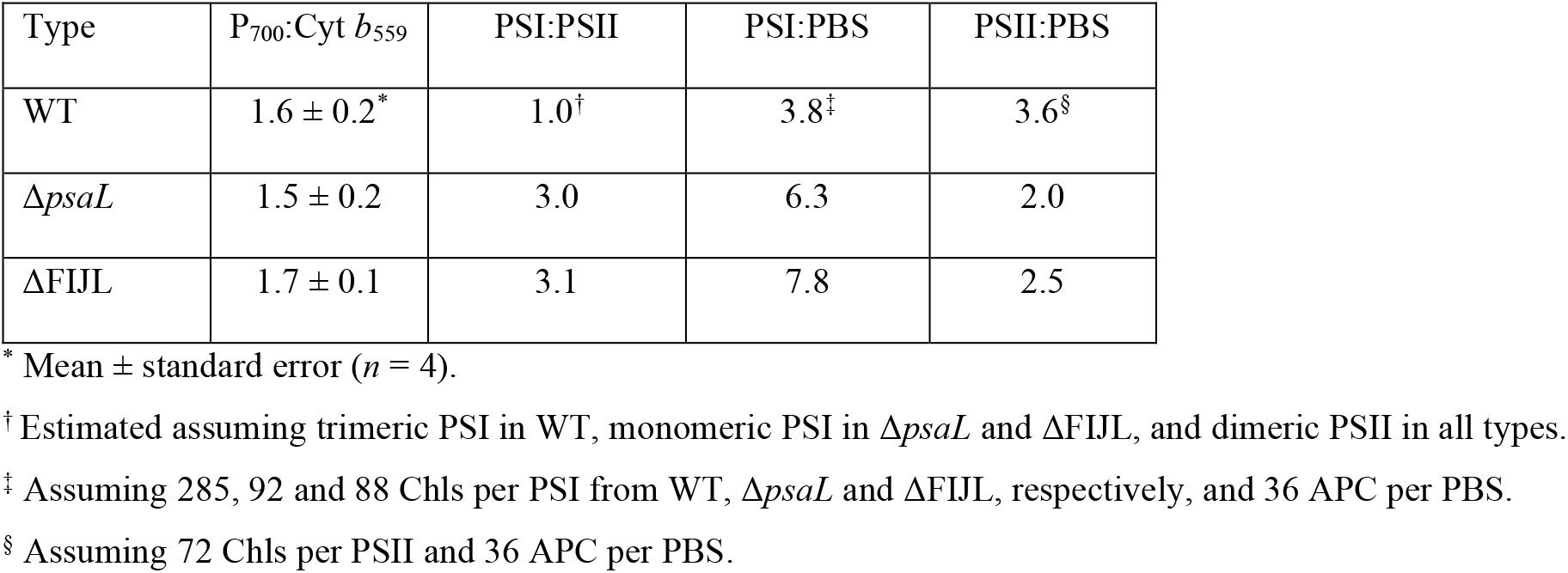
Stoichiometric ratios of PSI, PSII and PBS

| Type | P<sub>700</sub>:Cyt *b*<sub>559</sub> | PSI:PSII | PSI:PBS | PSII:PBS |
| --- | --- | --- | --- | --- |
| WT | 1.6 ± 0.2* | 1.0<sup>†</sup> | 3.8<sup>‡</sup> | 3.6<sup>§</sup> |
| *ΔpsaL* | 1.5 ± 0.2 | 3.0 | 6.3 | 2.0 |
| *ΔFIJL* | 1.7 ± 0.1 | 3.1 | 7.8 | 2.5 |

\* Mean ± standard error (*n* = 4).
<sup>†</sup> Estimated assuming trimeric PSI in WT, monomeric PSI in *ΔpsaL* and *ΔFIJL*, and dimeric PSII in all types.
<sup>‡</sup> Assuming 285, 92 and 88 Chls per PSI from WT, *ΔpsaL* and *ΔFIJL*, respectively, and 36 APC per PBS.
<sup>§</sup> Assuming 72 Chls per PSII and 36 APC per PBS.

### Steady-state fluorescence emission spectra

As a further confirmation of the pigment stoichiometry changes, we recorded fluorescence emission spectra of intact WT, Δ*psaL*, and ΔFIJL cells at 77 K (Figure 1). The spectra recorded with 580 nm excitation (primarily absorbed by PC) have peaks at 650, 660, 687/693, and 720 nm. The peaks at 650 and 660 nm correspond to PC and APC and the ones at 687/693 and 720 nm – primarily to PSII and PSI, respectively. In accordance with the higher amount of PC determined in the WT, the spectra showed significantly more intense emission at 650 nm. The two monomeric PSI types, Δ*psaL* and ΔFIJL, had similar fluorescence spectra. The ratio of fluorescence emitted at 650 nm to 660 nm decreased in the monomeric PSI types in line with the decreased PC:APC ratio. Statistically significant changes in the PC:APC emission ratio were also found in the room-temperature fluorescence emission spectra (Supplementary Fig. S3).

**Figure 1.**
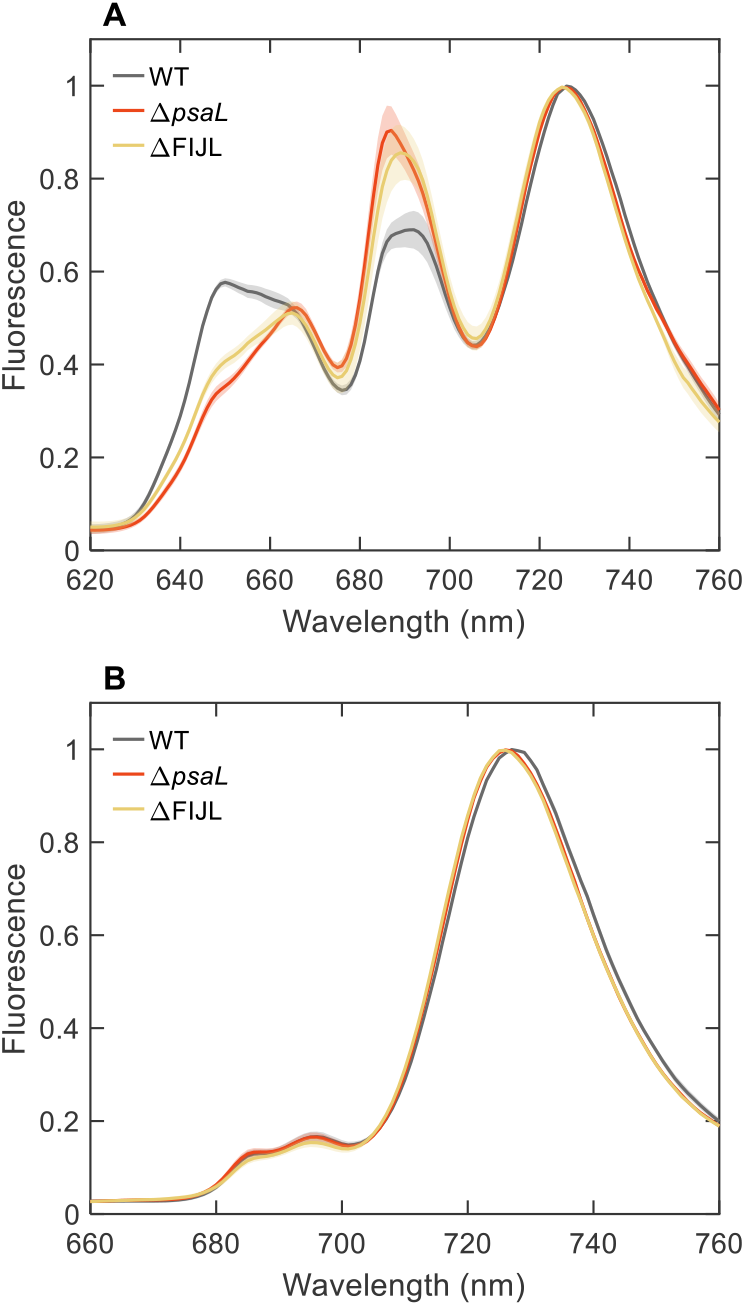
Fluorescence emission spectra of intact cells of *Synechocystis* WT, Δ*psaL* and ΔFIJL recorded at 77 K, normalized to the maximum at 720 nm. (A) Excitation wavelength 580 nm; (B) Excitation wavelength 440 nm. The spectra are average from 5–7 independent experiments; the shaded area represents the standard error.

The photon energy absorbed by the PBSs is ultimately distributed between both PSI and PSII as evident from their corresponding emission peaks. The relative amplitudes of the 687/693 and 720 nm peaks suggest that in Δ*psaL* and ΔFIJL cells more energy is transferred to PSII compared to WT. In contrast, the fluorescence spectra recorded with 440 nm excitation (almost exclusively absorbed by the photosystems) show no significant difference in the ratio of the PSII and PSI peaks. This is expected because the PSI:PSII stoichiometric ratio (on monomer basis) is unchanged. Thus, the stronger PSII emission upon 580 nm indicates that the distribution of excitation energy from PBSs to PSII/PSI is altered in the monomeric PSI types.

### P_700_ oxidation kinetics

To compare the effective antenna size of PSI, we recorded the oxidation kinetics of P_700_ (absorption transients at 830 nm) in intact cells and isolated PSI complexes upon illumination in the presence of DCMU and MV. DCMU prevents re-reduction of P_700_^+^ by electrons from PSII and MV accepts electrons from PSI keeping the acceptor side of PSI in oxidized state and minimizing cyclic electron flow. The oxidation curves at different light intensities are shown in Supplementary Fig. S4 and the oxidation rates obtained by fitting logistic or exponential kinetics to the curves are in Figure 2. At all light intensities in the range 6–805 μmol m^−2^ s^−1^, the oxidation rates in WT *Synechocystis* cells were higher than either of the monomeric PSI types – up to 60% at 140 μmol m^−2^ s^−1^. These data indicate that the effective antenna size of monomeric PSI in vivo is smaller compared to trimeric PSI. The oxidation kinetics were measured with 635 nm light, predominantly absorbed by the PBS. Hence, a possible explanation for the different rates could be that PBS transfer energy more effectively to trimeric than to monomeric PSI. This hypothesis is supported by the fact that the P_700_ oxidation rates were similar in monomeric and trimeric isolated PSI (Figure 2B) as well as in thylakoid membranes, which lack PBSs (Supplementary Fig. S5).

**Figure 2.**
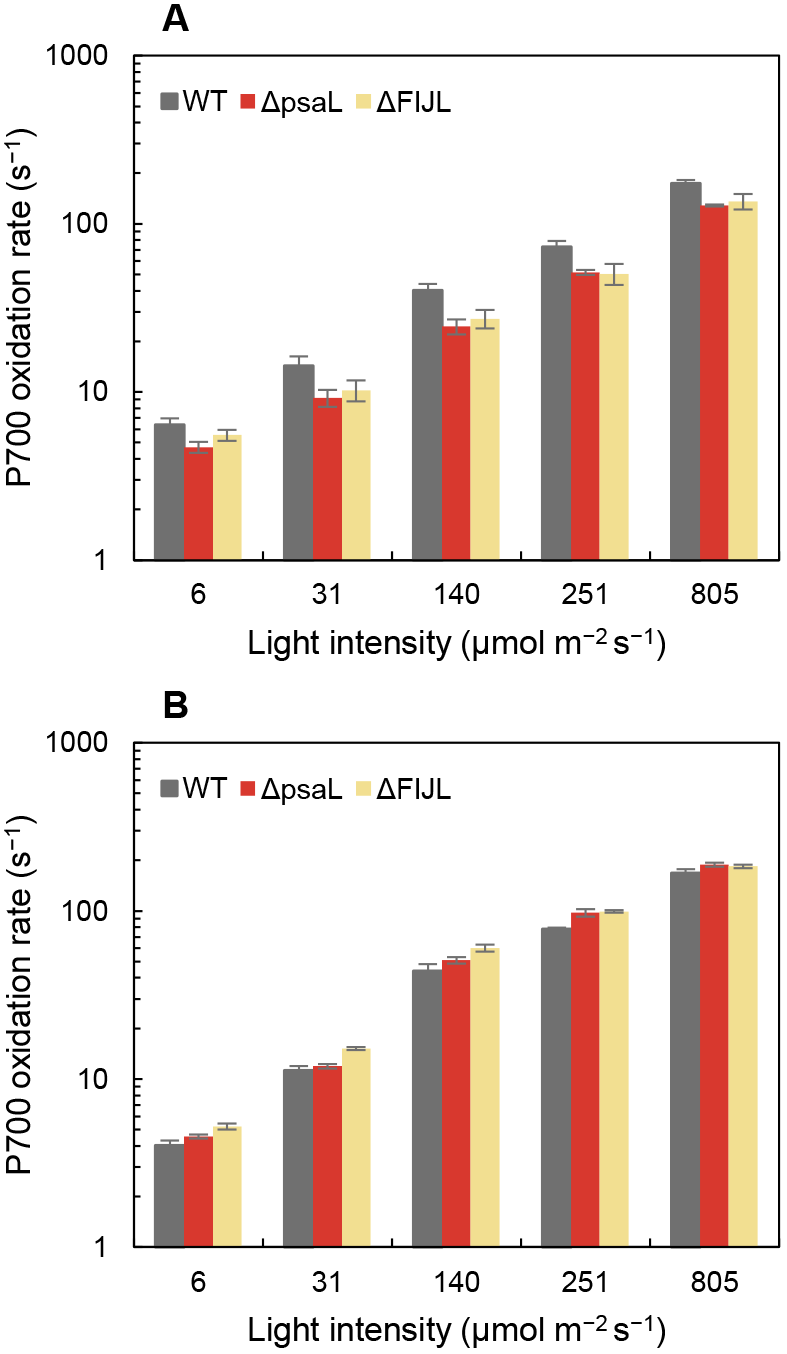
P_700_ oxidation rates. (A) intact cells of *Synechocystis* WT, Δ*psaL* and ΔFIJL; (B) isolated PSI. The rates are calculated from the differential absorption at 830 nm induced by pulses of different light intensity, applying logistic or exponential fit (for intensities under and above 100 μmol m^−2^ s^−1^, respectively). Error bars indicate standard errors from four independent experiments. Note the logarithmic vertical scale.

### Fluorescence kinetics of cells at room temperature

We employed picosecond time-resolve fluorescence spectroscopy to gain better understanding of the excitation energy partitioning in the intact cells of *Synechocystis* with trimeric (WT), monomeric (Δ*psaL*) and subunit-depleted (ΔFIJL) PSI. Time-resolved fluorescence enables better separation of emission components, for example emission of PBSs that are energetically coupled to the photosystems from free PBPs and can detect changes in the architecture and supramolecular organisation of the photosynthetic complexes that affect the EET. We applied global five-exponential analysis of the fluorescence decays recorded in the wavelength range 600–720 nm after excitation at 580 nm. Figure 3 compares the lifetimes and decay-associated emission spectra (DAES) of WT, Δ*psaL*, and ΔFIJL cells. The lifetimes and spectra are very similar between genotypes and are comparable to previously published results on cyanobacterial cells (Mullineaux and Holzwarth, 1991; Tian et al., 2011; Akhtar et al., 2020). The first two components (Figure 3A,B), with respective lifetimes around 30 ps and 90–100 ps, represent EET within the PBS (Akhtar et al., 2020). The DAES have characteristic positive and negative peaks around 650, 660, and 680 nm, signifying decay and rise of the emission from PC, APC and red-shifted APC (APC_680_), respectively. The third component with a lifetime about 170 ps represents mainly decay of APC excitations in PBS coupled to photosystems and the fourth, around 500–600 is associated with trapping in PSII. A longer-lived component (1.4–1.6 ns, not plotted) indicated the presence of a negligible amount (2–3%) of uncoupled PBPs.

**Figure 3.**
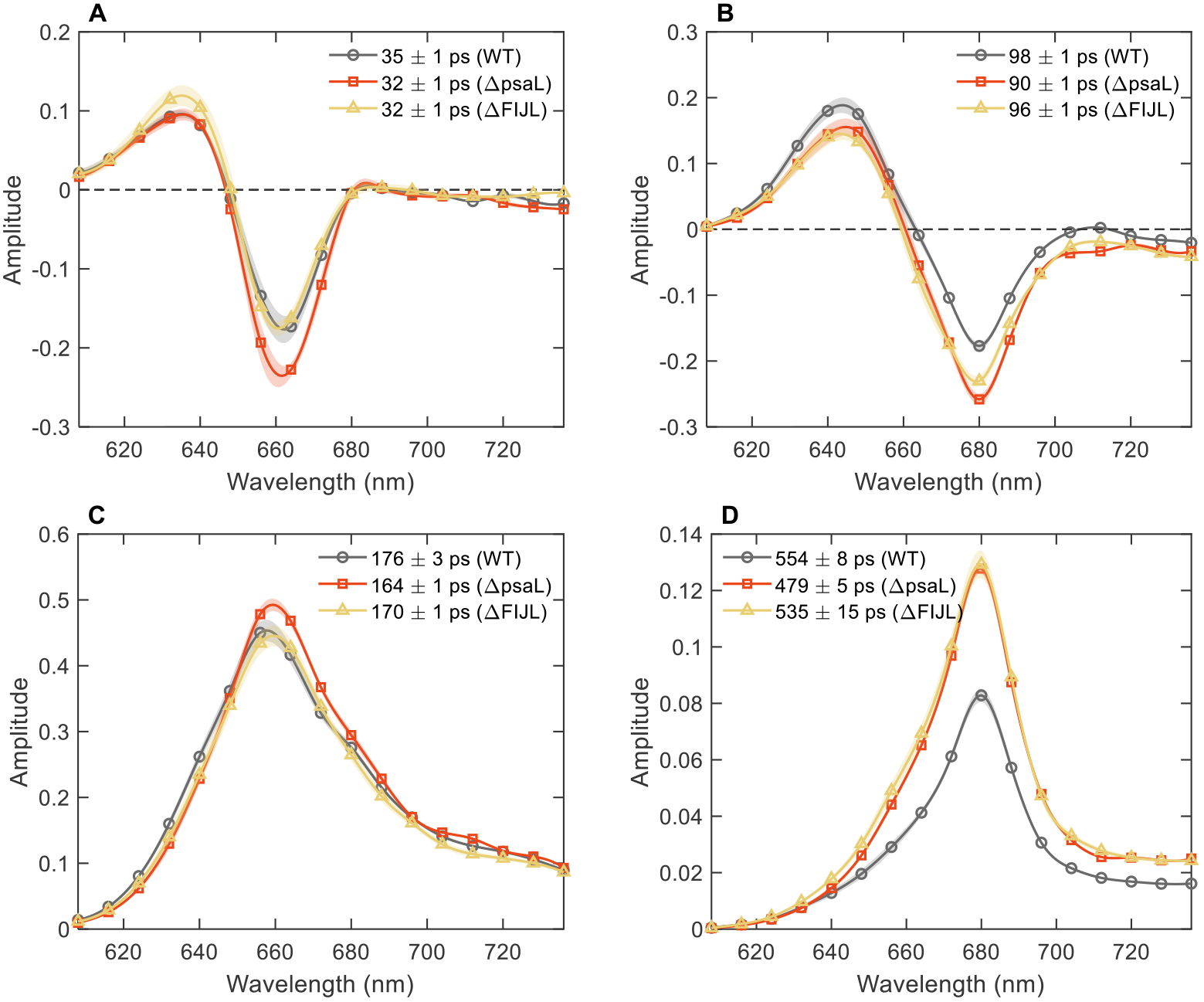
DAES of *Synechocystis* cells obtained by global lifetime analysis of the fluorescence decays recorded at room temperature with 580 nm excitation. Panels A-D compare individual DAES for WT (black circles), Δ*psaL* (red squares), and ΔFIJL (gold triangles) cells. The longest-lived (1.3-1.6 ns) component is omitted. The spectra are average from six independent experiments, normalized to the same integrated fluorescence intensity (area of the steady-state fluorescence spectrum). The standard error of the mean is shown by the shaded areas.

As seen in Figure 3, there is little difference between the genotypes as concerning the shape of the DAES. On close inspection, the DAES of WT cells display relatively larger amplitudes at 640–650 nm (Figure 3B,C), consistent with their higher PC:APC ratio. There was also a small increase in the PBS-associated decay lifetimes in WT compared to the other two types.

The most notable kinetic difference between WT cells and those with monomeric PSI is in the PSII decay component (Figure 3D). In monomeric PSI types, the amplitude of this component was 50±5% larger than in WT (the difference is statistically significant at α = 0.01). This result confirms the finding that a larger proportion of excitation energy is transferred to PSII in cells with monomeric PSI, compared to cells with trimeric PSI. Moreover, as we observe only negligible amount of uncoupled PBSs in the *Synechocystis* cells, we can make the reverse conclusion – namely, that PBSs transfer more energy to PSI when it is trimeric rather than monomeric.

### Fluorescence kinetics of cells at 77 K

We further examined the fluorescence decay kinetics of intact cells at 77 K for better resolution of the different pigment groups. Most importantly, the emission from the long-wavelength “red” Chls in PSI is well pronounced allowing us direct comparison of the energy flow to the two photosystems. Global analysis of the fluorescence decays resulted in six decay lifetimes and DAES. Individual DAES for the three *Synechocystis* genotypes are compared in Figure 4. We will first briefly describe the kinetic components in WT cells. The fastest component (13 ps, Figure 4A) shows decay of PC emission at 640 nm and concomitant rise of APC emission at 660 nm. In addition, EET from bulk to red Chls in PSI occurs on this timescale (690 to 720 nm). The second component (56 ps, Figure 4B) shows decay of both PC and APC and rise of the red-shifted APC_680_. The 145-ps component (Figure 4C) has positive peaks corresponding to all three PBP groups (PC, APC and APC_680_) that evidently decay as energy is transferred to Chls. Also notable are the negative peaks at 690 nm (PSII) and 720 nm (PSI). The PSII and PSI emission components decay mainly with lifetimes of 373 ps and 982 ps (Figure 4D,E) while the final component (3.8 ns, Figure 4F) is of negligible amplitude. Remarkably, PC fluorescence detected at 640–650 nm spans the entire range of decay timescales (in WT). A sizeable fraction of long-lived PC is less efficient in transferring excitation energy downstream (note the 650 nm peaks in Figure 4C–E). It is also of note that a considerable proportion of photon energy absorbed by the PBS is delivered to PSI, judging by the height of the DAES peaks around 720 nm.

**Figure 4.**
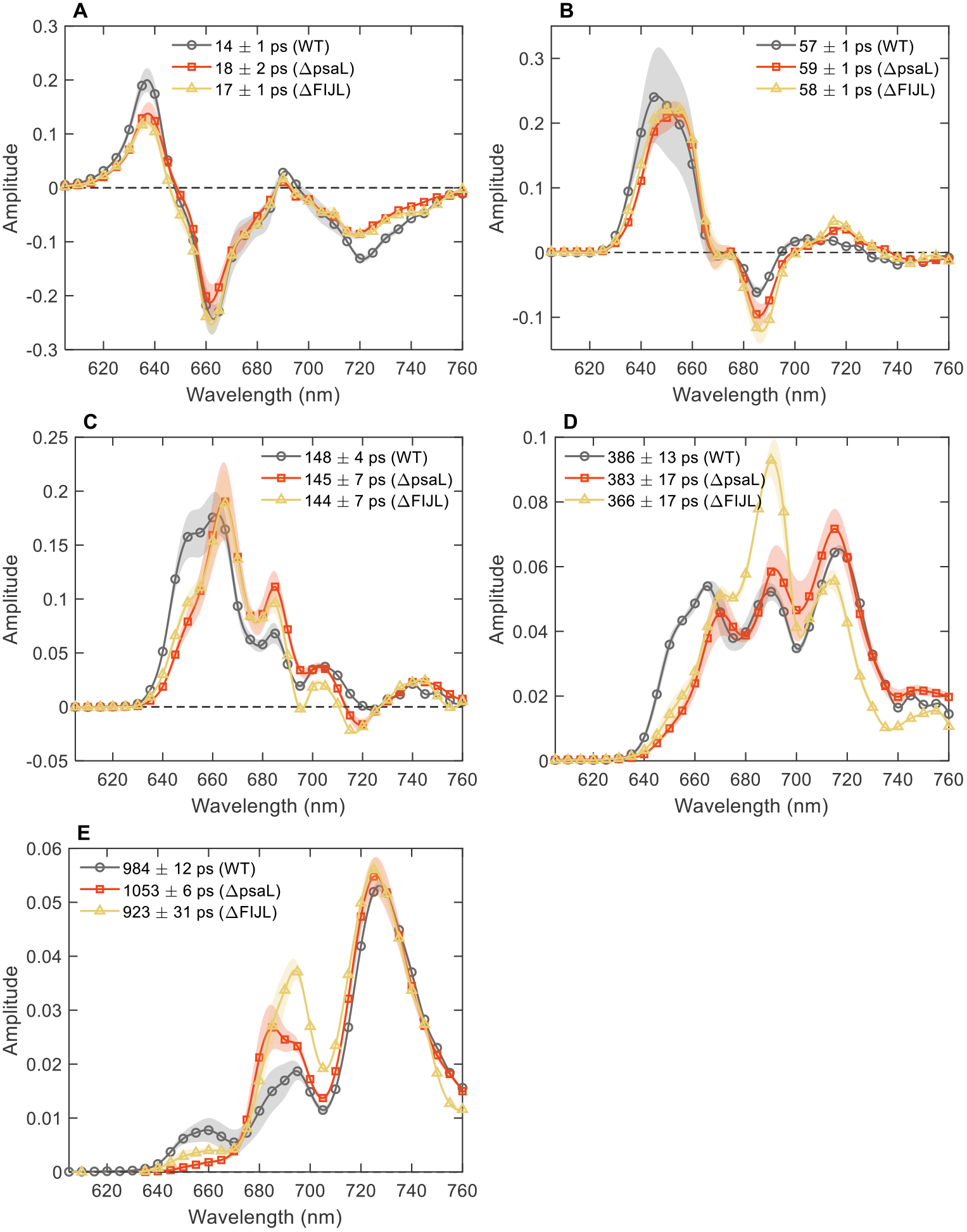
DAES of *Synechocystis* cells obtained by global lifetime analysis of the fluorescence decays recorded at 77 K with 580 nm excitation. The data are normalized to the same integrated fluorescence intensity. Panels A-F compare individual DAES for WT (black circles), Δ*psaL* (red squares), and ΔFIJL (gold triangles) cells. The spectra are averages of three independent and one technical replicate. The shaded areas represent standard error. A small-amplitude component with 2–3 ns lifetime is omitted.

Compared to WT, the DAES of both monomeric PSI types showed notable differences. Remarkable are the kinetic differences in the 640–680 nm wavelength range reflecting PBPs emission. In Δ*psaL* and ΔFIJL cells, PC emission decays faster than in WT – the peaks at 650 nm in the long-lived DAES all but vanish. Conversely, the amplitudes of the APC peaks at 660 and 680 nm are larger than in WT. Especially obvious is the larger negative peak at 680 nm in the 58-ps DAES (Figure 4B) and corresponding positive peak in the 136-ps DAES (Figure 4C). These results show unequivocally that the altered PBP stoichiometry results in changes in the excitation dynamics of the intact and energetically coupled PBSs.

The fluorescence emission components decaying with lifetimes around 140 ps and 360 ps (Figure 4C-D) have higher amplitudes at 685 nm in the monomeric PSI types. Although the emission at 685 nm can originate from the terminal emitters in APC (APC_680_), the decay is nevertheless due to energy being transferred to PSII. These components are absent in PSII-deficient *Synechocystis* (unpublished data). Therefore, the fluorescence kinetics at 77 K further supports the finding that a larger proportion of PBS excitations are transferred to PSII when PSI is monomeric.

As a control, we analysed the fluorescence kinetics at 77 K upon 440 nm excitation, which excludes the PBS contribution to the dynamics. In this case, the lifetimes and DAES of intact cells were virtually identical to the ones reported earlier for isolated thylakoid membranes (Supplementary Fig. S6). The fluorescence decays reflect mainly the dynamics in PSI, as it has 4–5 times larger absorption cross section than PSII. The most apparent difference in the fluorescence kinetics is the reduced amplitude of the DAES emitting at 710–715 nm, attributed to the loss of red Chls at the trimerization region (Akhtar et al., 2021).

Additionally, we compared the fluorescence kinetics of isolated PBSs of WT, Δ*psaL*, and ΔFIJL at room temperature and 77 K (Supplementary Figs. S7, S8). Interestingly, the clear differences in the PC dynamics, that were observed in whole cells, could not be detected in the isolated PBS. This result is in line with the fact that changes in the PC:APC ratio were detected in whole cells but not in isolated PBSs.

## DISCUSSION

A key finding of this study is that the oligomeric state of PSI exerts control over the abundance of PBPs in *Synechocystis* and, by extension, the PBS composition. The Δ*psaL* and ΔFIJL genotypes with monomeric PSI contained more PBS cores on Chl basis compared to the WT genotype with trimeric PSI but, at the same time, fewer PCs per APC core (Tables 1 and 2). In agreement with the spectroscopic quantification, semiquantitative SDS-PAGE analysis showed lower PC:APC ratios in cells of the monomeric PSI mutants (Supplementary Fig. S9). The results are in line with the reported fluorescence spectral changes in the Δ*psaL* mutant of *Synechocystis* (Kłodawska et al., 2020).

By comparing the data from cells and isolated PBSs we can conclude that WT cells contain extra PC rods that are not found in the isolated PBS. There are two possibilities to consider. The first is that the additional PC are weakly bound to the PBS making up for longer rods radiating from the PBS core or additional laterally bound rods that are dissociated during the PBS isolation. In support of the longer-rod hypothesis, we found a change in content of the two rod linkers, L_R_^33^ (CpcC1) and L_R_^30^ (CpcC2), in the SDS-PAGE of cell soluble material (Supplementary Fig. S9). In both monomeric PSI types the abundance of L_R_^30^ relative to L_R_^33^ was significantly lower than in WT. Assuming a model, where L_R_^30^ connects hexamers distal to the APC core, whereas L_R_^33^ is bound to the proximal hexamers (Ughy and Ajlani, 2004), the results can be interpreted to show that some PBS in the monomeric PSI types have rods composed of only two hexamers.

Larger PBS should result in increased rod-core energy equilibration times (Sandström et al., 1988; Zhang et al., 1997). Although the energy equilibration was indeed longer in WT compared to the other two types, the differences are too small to account for an almost double PC content per core. On the other hand, a very long-lived PC fluorescence component in WT can be assigned to a fraction of weakly connected PC rods that suggest unconventional attachment to the membrane. These “semi-free” rods may be easily lost during PBS isolation.

The second possibility is that the “extra” PC in WT cells is assembled in distinct PBSs that do not contain APC. It is tempting to assign this to the PBS type containing the linker protein CpcG2 or CpcL. CpcL-containing PBS (CpcL-PBS) consist only of PC rods attached directly to the membrane without an APC core and are known to transfer energy preferentially to PSI (Kondo et al., 2007; Kondo et al., 2009). CpcL-PBS are readily isolated from APC-deficient cyanobacterial mutants, where they become the predominant PBS type but their abundance in WT *Synechocystis* cells is not well documented and appears to vary with experimental conditions and especially light quality, as the expression of CpcG2/CpcL is regulated by the phytochrome-like protein CcaS (Hirose et al., 2008). SDS-PAGE showed no major differences in the PBPs or linker protein content of the isolated APC-containing PBSs (Supplementary Fig. S9), nor were there notable functional differences among them. It is possible that the reduced PC:APC ratio in both mutants with monomeric PSI is because these mutants, in contrast to WT, do not assemble CpcL-PBSs. In this case it would follow that almost half of the PC in WT is assembled in CpcL-PBSs (comparing the PC:APC ratios of cells and isolated PBSs). It is worth pointing out that the observed reduction in the ratio of L_R_^30^/L_R_^33^ linkers in the monomeric PSI types is also compatible with a lower amount of CpcL-PBSs since these PBSs contain less CpcC2 (L_R_^30^) compared to the CpcG-type PBSs (Liu et al., 2019).

The modulation of the PBS composition (or architecture) can be understood as an adaptive response. Cyanobacteria regulate their PBS content in response to the physiological conditions. Growth under high irradiance changes the abundance of PBS, reducing the cell PC content (Raps et al., 1985) and shortening of the PC rods has been reported in *Synechococcus* (Samson et al., 1994), *S. maxima* (Garnier et al., 1994), and *S. platensis* (Nomsawai et al., 1999). The growth conditions regulate the expression of specific linker proteins controlling the PBS architecture (Nomsawai et al., 1999; Hihara et al., 2001), such as the CpcG2/CpcL linker (Hirose et al., 2008). PBPs are also reservoirs for nutrients, degraded when nutrients, e.g. nitrogen, are scarce (Salomon et al., 2013), so it is plausible that downstream metabolism changes will also alter the PBP content.

We reason that the absence of trimeric PSI in *Synechocystis* results in suboptimal or imbalanced EET, consequently electron transfer, which causes a change in the PBS content and composition. The steady-state and time-resolved fluorescence data show that more energy is transferred to PSII in WT and that the effective PSI antenna size is diminished in Δ*psaL* and ΔFIJL cells (but not thylakoid membranes or isolated PSI). The results strongly suggest that the PBSs transfer excitation energy more efficiently to trimeric than to monomeric PSI. There could be several reasons for this difference:

1. Supercomplexes of PBSs with trimeric PSI can be more efficient by sharing the PBS among all three PSI RCs – increasing the effective antenna size. Then the increased number of APC-PBSs in the monomeric PSI strains can be seen as compensatory response for the loss of effective absorption cross-section. However, as PBSs transfer energy to both photosystems (Ashby and Mullineaux, 1999), merely increasing the number of PBSs will exacerbate the energy imbalance rather than alleviate it. On the other hand, strains devoid of PBSs or with truncated PBSs rods compensate for the reduced PSII antenna size by increasing the abundance of PSII (Ajlani et al., 1995; Nagarajan et al., 2014; Liberton et al., 2017) or the number of PBSs per photosystem (Leganes et al., 2014).
2. PSI trimers might have higher affinity to the PBSs forming a more stable PBS-PSI supercomplex. This may be in contrast to structural modelling that suggests the existence of a PBS fraction directly attached to PSI monomers (Zlenko et al., 2016).
3. CpcL-PBS may be an efficient antenna of trimeric PSI (in WT) but not of monomeric PSI. At present, this is only a conjecture based on the hypothetical assignment of the additional PC rods found only in WT cells to CpcL-PBS. It can be postulated that CpcL-PBS are only stably assembled in the presence of trimeric PSI. In *Anabaena sp.* PCC 7120, CpcL-PBS has been shown to attach to the tetrameric PSI complex at the interface between two protomers (Watanabe et al., 2014), effectively transferring energy to the PSI core within ~90 ps (Noji et al., 2021). It could be hypothesized that in *Synechocystis* CpcL-PBS is only stably attached to two protomers of the PSI trimer. This hypothesis would need further verification, firstly to confirm that the CpcL-PBS content differs in cells with monomeric and trimeric PSI, and then, to explore the affinity and coupling of CpcL-PBS to PSI.
4. We must also consider the role of state transitions (Mullineaux and Emlyn-Jones, 2005; Calzadilla and Kirilovsky, 2020). The Δ*psaL* mutant of *Synechococcus* PCC 7002 was found to be capable of performing a state II–state I transition faster than the wild type (Schluchter et al., 1996; Aspinwall et al., 2004). However, there is no indication that the mutant is locked or preferentially found in state I. On the contrary, as state transitions are regulated by the redox state of the PQ pool (Mullineaux and Emlyn-Jones, 2005), it should be expected that any relative loss of PSI excitation will shift the balance toward state II (which would cause an opposite change in the fluorescence spectra than observed).

It is also possible that a combination of the above factors contributes to the changes in the excitation energy distribution. Regardless of the mechanism, however, the results show that the oligomerization state of PSI has a significant impact on the excitation energy flow from PBSs to the photosystems in *Synechocystis*. Mutants with monomeric PSI compensate for the imbalanced excitation by adjusting the PBS composition. In contrast, we could not discern a particular role of PsaF in the EET from PBS to PSI as previously suggested (Fromme et al., 2003). These results add to the existing body of evidence that the PBSs are a remarkably responsive and tuneable light-harvesting antenna system but also provide a hint of the evolutionary advantage of oligomeric PSI in conjunction with PBSs.

## Acknowledgements

We are grateful to Prof. Nathan Nelson and Prof. Dario Leister for providing us the ΔFIJL strain of *Synechocystis.* We thank Reviewer 1 for valuable suggestions regarding the discussion and interpretation of the results.

---

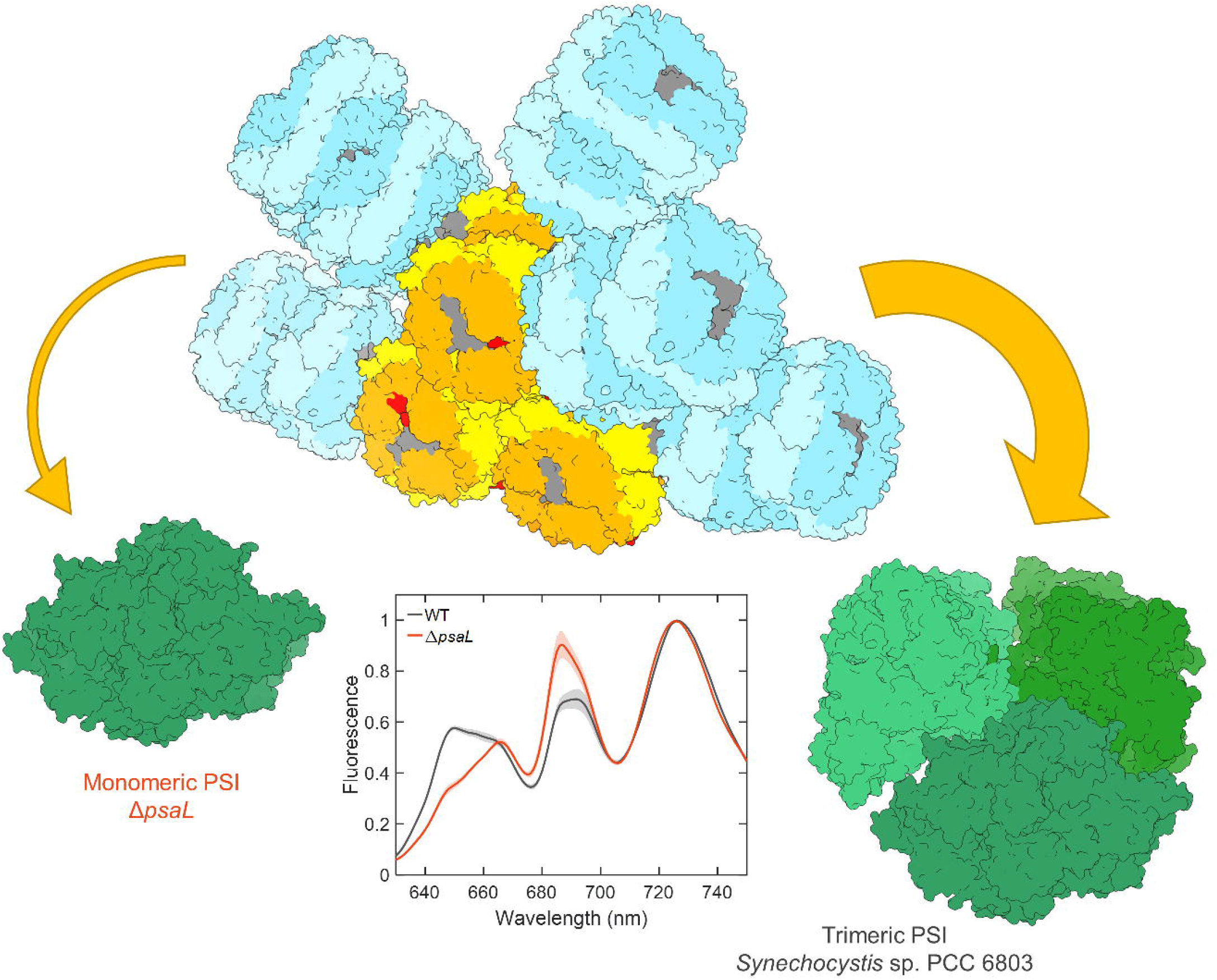

