## Supplementary Material for "Trimeric Photosystem I facilitates energy transfer from phycobilisomes in *Synechocystis* sp. PCC 6803"

### SUPPLEMENTARY FIGURES

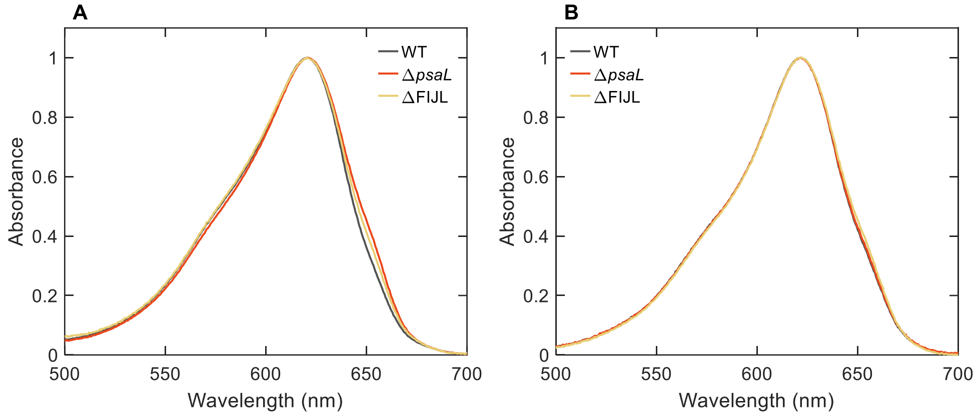


Figure S 1. Absorption spectra of (A) total phycobiliprotein extract of the cells; (B) isolated phycobilisomes. Spectra are normalized to the maximum.

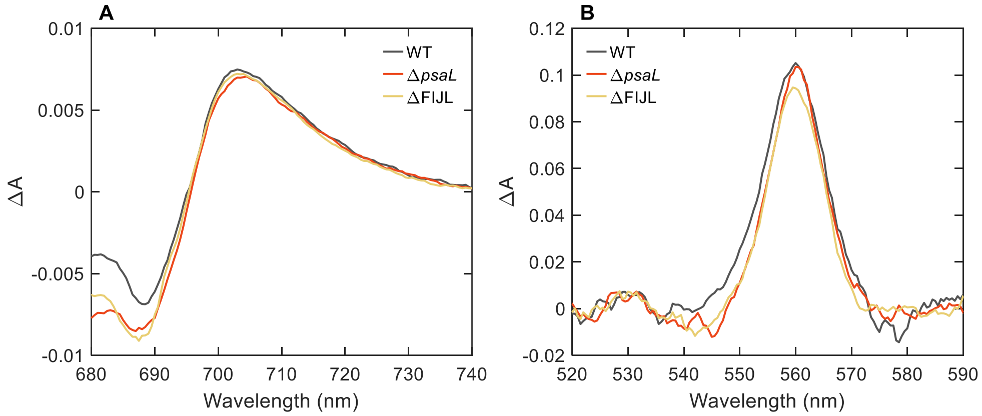


Figure S 2. Spectrophotometric determination of $P_{700}$ and Cyt $b_{559}$. (A) Absorption difference spectra of $P_{700}$ in thylakoid membranes of *Synechocystis* – ascorbate-reduced minus ferricyanide-oxidized samples. (B) Absorption difference spectra – ascorbate-reduced minus hydroquinone-reduced samples.

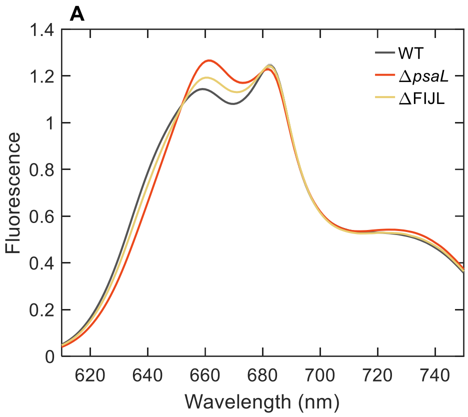


Figure S 3. Steady-state room-temperature fluorescence emission spectra of WT and mutant cell suspensions. (A) 580 nm light exciting mainly PC in PBS; (B) 440 nm light exciting mainly chlorophylls in photosystems. The spectra, normalized to the same area, are average of 12 independent experiments. The shaded areas represent standard error.

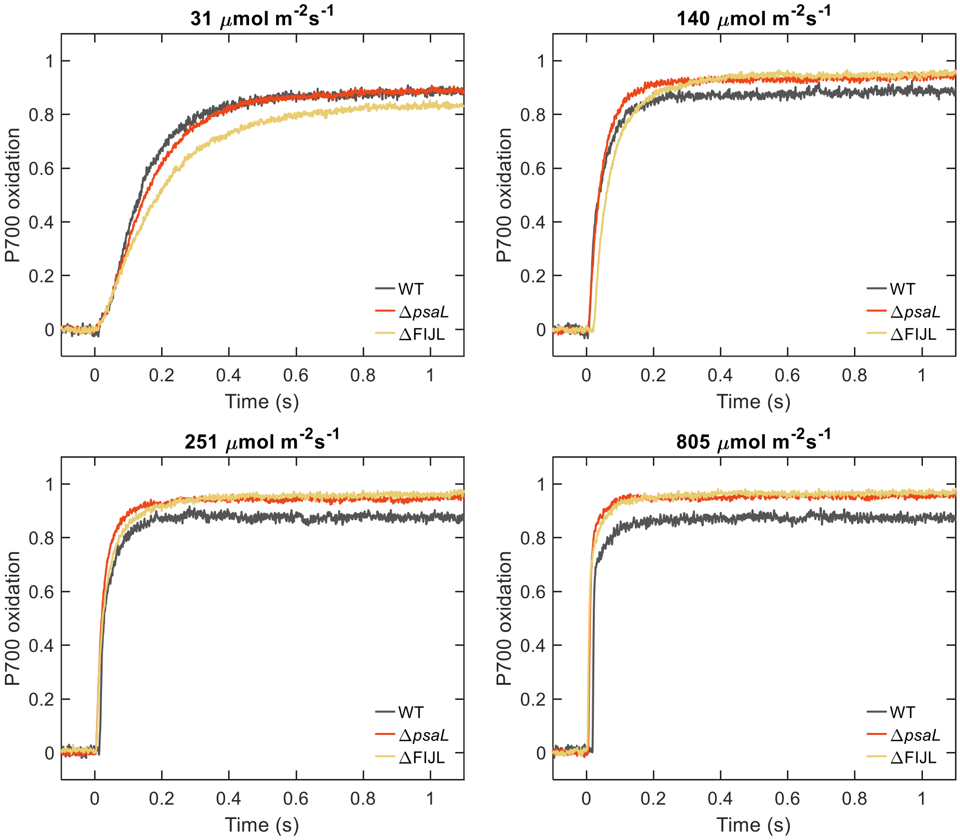


Figure S 4. Typical $P_{700}$ signals of *Synechocystis* WT, Δ*psaL* and ΔFIJL cells treated with MV and DCMU. Actinic light intensity is indicated on top of the plots. Signals are normalized to 1 at maximum.

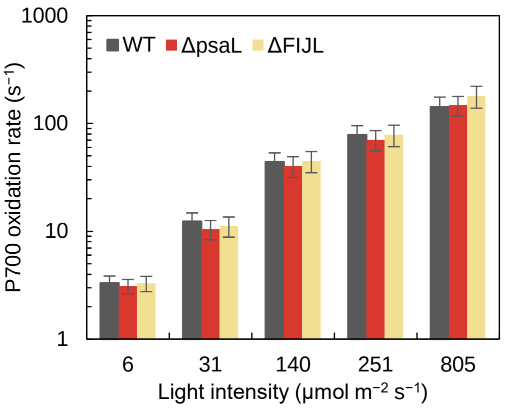


Figure S 5. $P_{700}$ oxidation rates in isolated thylakoid membranes of *Synechocystis* WT, Δ*psaL* and ΔFIJL. Error bars indicate standard errors from four independent experiments.

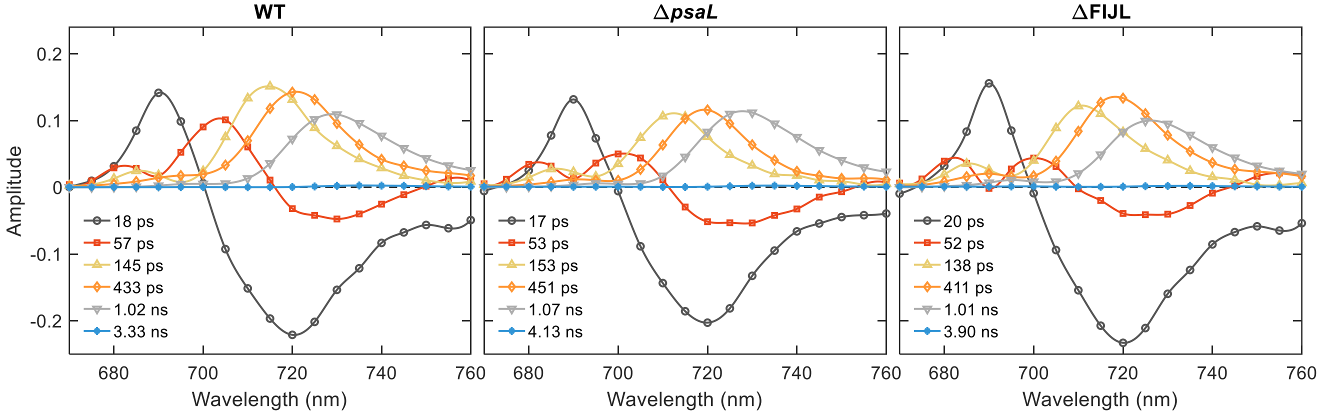


Figure S 6. Decay-associated fluorescence emission spectra of intact cells of *Synechocystis* (WT, Δ*psaL* and ΔFIJL mutants) obtained from global analysis of fluorescence decays measured at 77K upon 440 nm excitation. The spectra are scaled to the same spectrally-integrated fluorescence intensity.

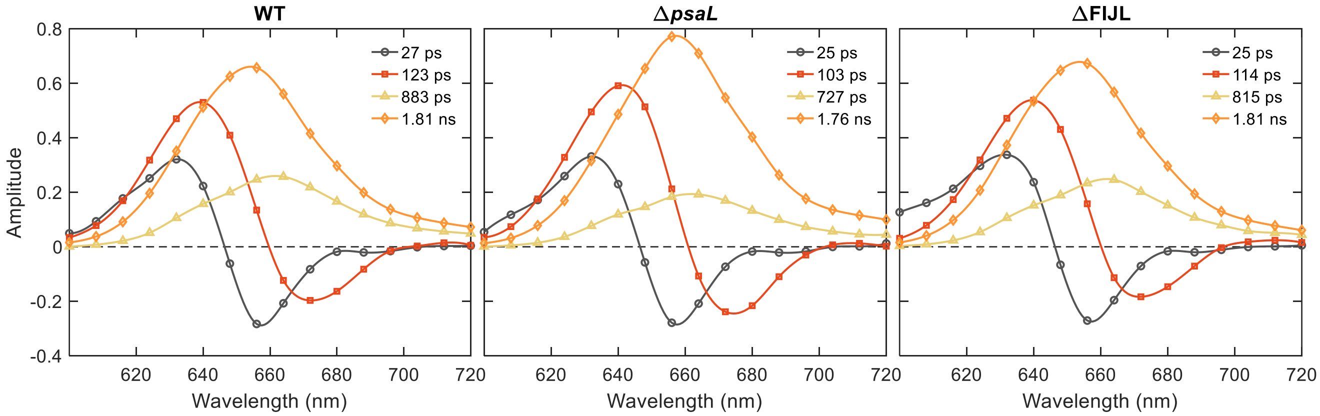


Figure S 7. Decay-associated fluorescence emission spectra of isolated PBS of *Synechocystis* (WT, Δ*psaL* and ΔFIJL mutants) obtained by global lifetime analysis of the fluorescence decays recorded at RT with 580 nm excitation. The spectra are scaled to the same spectrally-integrated fluorescence intensity.

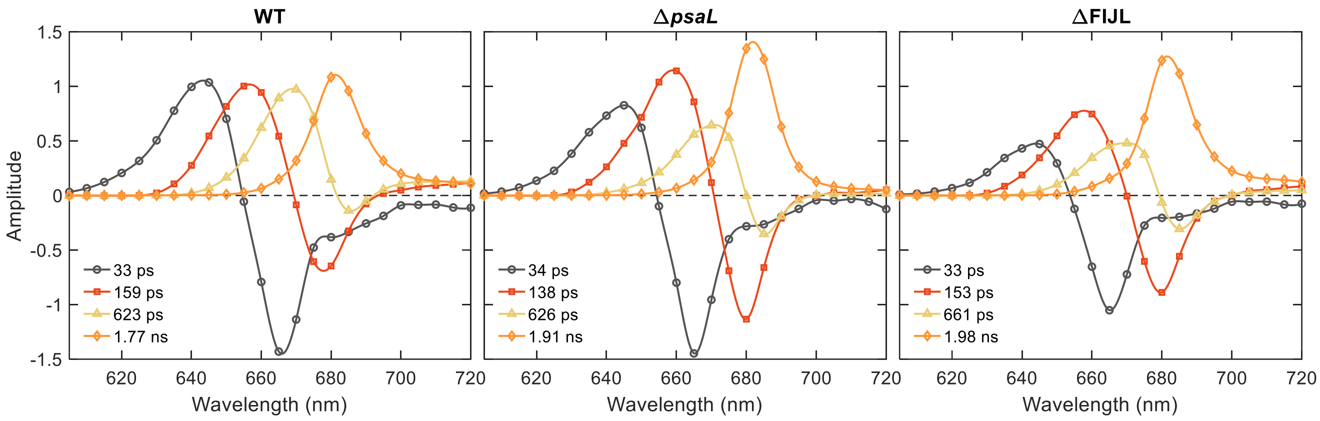


Figure S 8. Decay-associated fluorescence emission spectra of isolated PBS of *Synechocystis* (WT, Δ*psaL* and ΔFIJL mutants) obtained by global lifetime analysis of the fluorescence decays recorded at 77K with 580 nm excitation. The spectra are scaled to the same spectrally-integrated fluorescence intensity.

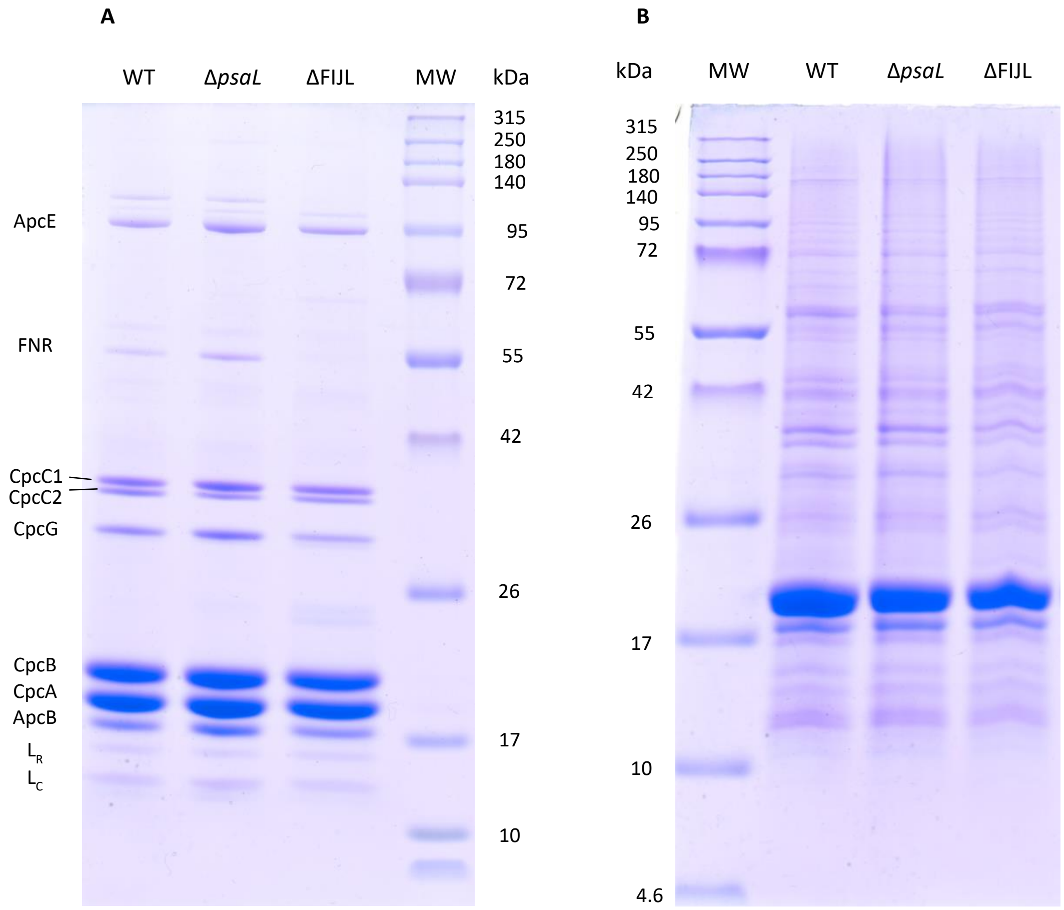


| | WT | Δ*psaL* | ΔFIJL |
|---|---|---|---|
| PC:APC | **6.8** | **5.7** | **7.3** |
| CpcC1:CpcC2 | **1.7** | **3.4** | **2.4** |

| | WT | Δ*psaL* | ΔFIJL |
|---|---|---|---|
| PC:APC | **5.9** | **4.5** | **4.9** |
| CpcC1:CpcC2 | **1.6** ±0.1 | **2.5** ±0.2 | **2.1** ±0.1 |

Figure S 9. Tricine-SDS-PAGE of isolated PBSs (A) and soluble cell extracts (B) from *Synechocystis* WT, Δ*psaL* and ΔFIJL. Protein composition was determined by Tricine-SDS-PAGE (Hermann Schägger, Nature Protocols 2016, 1, 16−22). Equal amounts of proteins were denatured at 85°C for 5 minutes, separated on a 10–16% gradient gel and then stained by standard Coomassie R-250 staining protocol. Numbers below the gel images show the ratio of the CpcA+CpcB band intensity to ApcB and CpcC1 to CpcC2. The CpcC1:CpcC2 ratios in (B) are mean ± SE ($n$ = 5). Densitometric analysis was done using GelAnalyzer 19.1 (www.gelanalyzer.com) by Istvan Lazar Jr., PhD and Istvan Lazar Sr., PhD, CSc.

### SUPPLEMENTARY TABLES

Table S 1 ratios of PC to APC in the isolated PBSs calculated from the absorption spectra

| Type | PC:APC |
|---|---|
| WT | 3.8 ± 0.1 |
| ΔpsaL | 4.2 ± 0.1 |
| ΔFIJL | 3.6 ± 0.3 |
